# “High resolution discovery of regulatory DNA with synthetic wild-type and ablated genome constructs”

**DOI:** 10.1101/2020.11.02.355818

**Authors:** Benjamin Holmes, Lin Lin, Richard Sherwood, David Gifford

## Abstract

Genomic enhancer elements play a role in modulating gene expression by interacting with effector proteins known as transcription factors. We present a new computational method, “STARR-scan”, which identifies transcription factor motif appearances in synthetic DNA constructs that are correlated with enhancer activity. Using a tiled STARR-seq assay, we demonstrate that STARR-scan can suggest the biological function of transcription factors previously suspected to activate the APOBEC-3B gene. Using the same methodology, STARR-scan can suggest transcription factors with previously unknown biological activity which may regulate the APOBEC-3B gene. These novel factors may have biological significance in our understanding of cancer biology.

## INTRODUCTION

This work introduces STARR-scan, a method for enhancer analysis which discovers putative transcription factor motifs using the experimental behavior of synthetic DNA constructs in regulating mRNA production. The synthetic constructs are drawn from selected regions of the human genome, allowing us to suggest the regulatory potential of the discovered motifs. .

Our approach builds on previous methodologies for implicating transcription factors in gene expression through motif analysis. Whereas current approaches to motif discovery either look at small regions and miss potential interactions, or scan broad sequence regions and are limited in statistical and spatial resolution, the goal of this work is to facilitate high-resolution discovery of condition- and cell type-specific regulatory interactions using perturbative empirical analysis.

Using STARRScan, we demonstrate high-confidence discovery of specific transcription factor binding motifs in a genomic region that influence mRNA production in an extrachomosomal context. We demonstrate this method by running STARRscan to pinpoint apparent functional DNA sequence motifs which may implicate novel regulatory factors controlling the expression of the cancer-related APOBEC3B gene.

## BACKGROUND

Regulatory element DNA sequences in our genome, including promoters and enhancers, serve as part of the regulatory machinery that provides precise instructions for the correct expression of our genes. Understanding the identity and function of these regulatory elements is key for understanding human biology both in health and in disease. Promoters are located immediately upstream of the coding sequence of a gene they regulate and are typically easily identified. Enhancers can implement a complex cell-state specific regulatory program for a gene, and typically there are many enhancers for a human gene at varying genomic distances from the gene’s promoter. Identifying all of the enhancers that are active in a given cell state is an open problem in computational biology. This is in part because enhancers are bound by a wide variety of protein transcription factors that regulate their activity in a cell state specific manner. Thus characterizing the function of enhancers also involves discovering all of their cognate binding motifs for transcription factors. The full elucidation of promoters and enhancers is an important first step in understanding the internal programs of cells.

### Previous Work

The identification of active regulatory motifs and their associated transcription factors is a challenging problem that requires both experimental and computational methods that perturb genomic sequence to identify causal regulatory relationships. Disparate transcription factors can share the same DNA binding motif, and thus it is not always possible to identify the regulatory factors that may bind to an enhancer by using regulatory element sequence information. Identifying where and when identified transcription factors bind can be accomplished with methodologies such as ChIP-seq and ChIP-pet^1^. However, ChIP binding does not strictly imply function, as neutral binding is observed that has no regulatory consequence.

Perturbational screens of the genome with methods such as MERA^2^ can yield insights into DNA sequence regulatory elements in their native context, ^3^. These screens, permit regions of regulatory importance can be identified. Contemporary perturbational screens are limited in spatial resolution, and identify regulatory interactions with precision on the order of hundreds of bases. ChIP seq allows the experimenter to refine a search area and strengthen discovery confidence of computational methods once a target transcription factor has been identified. However, since ChIP-seq identified binding may be neutral, all ChiP-seq bound genomic regions must be validated for regulatory function using perturbation studies.

Contemporary genome cutting-based perturbational assays have their spatial resolution limited by three factors: (a) the uncertainty of the repair genotypes generated by a given gRNA induced genome cut, (b) the spatial limitations on where guide RNAs may be placed.One approach for analyzing perturbational assays is to gain statistical strength by binning results for gRNAs within a given genome sequence window. Using a larger bin size permits more reads to be used as evidence for a single bin, but reduces spatial precision of the data, creating a tradeoff between statistical strength of the evidence supporting a regulatory versus spatial precision. Frequently the binning resolution of gRNA based studies lends itself to the identification of enhancer areas on the order of tens or hundreds of bases, precluding confident resolution of motif elements which are typically associated with Transcription Factor binding consensus sequences spanning <30 nucleotides.

### Scope Of This Work

We use STARRscan to create confident predictions of specific, functional enhancer motifs using the analysis of the regulation of mRNA production by synthetic constructs. The use of designed constructs allows us to causally associate gene expression levels at high spatial resolution with DNA enhancer content. STARRscan uses information from expression levels of a large set of tiled oligos to propose high-confidence regulatory interactions which can be confidently associated with specific regulatory motifs that are located in a precise location within primary DNA sequences of the genome.

STARRseq overcomes the spatial resolution limitations of genome cutting methods by synthesizing a large number of computationally designed genomic variants. We do not rely upon the stochastic process of genome repair, using a dense CRISPR guide library attacking variously spaced enhancer loci for each candidate regulatory subregion. Instead we query the enhancer activity activity of computationally designed wild-type and systematically ablated DNA sequences containing or ablating 30 base pair oligo query sequences.

STARscan systematically queries putative enhancer sequences surrounded by contextual flanking sequences, and is a general approach to quantifying enhancer activity of sequences under study. Because it includes proximal native context, it is sensitive to limited context-dependent combinatorial effects. We targeted an understanding of the activation strength of ten to twenty base transcription factor motifs in enhancers, seeking to identify transcriptionally active sequence elements from an unbiased scan, and quantify their activation strength by comparing their differential activity with otherwise identical oligos having deletions of those sequences.

Our experimental design addressed (a) which sequence variants to test (b) using DNA synthesis to test perturbed and unperturbed genomic regions for enhancer activity and (c) multiplexed experiment design to provide evidence for activity at a given locus. We show how our experimental designcan be applied to infer a regulatory network for the APOBEC3B gene, and identify novel regulatory elements in this gene region, proposing interesting new regulatory interactions active in the cell in addition to specific sequence motifs which mediate these interactions and could provide targets for therapeutic interventions.

## METHODS

### Tiled oligo design, DNA barcodes, and RNA UMIs

We evaluated the regulatory potential of naturally-occurring DNA sequence motifs by designing a library of 150bp oligos tiling 91 regions identified in a CRISPR screen within a 600kb region around the APOBEC3B gene. Within each of the 91 regions, a series of 150 base pair “wild-type” oligos were chosen using a stride of 30 bases to tile each of the 91 regions. A total of 2000 wild type oligos were synthesized.

From each wild-type oligo five additional synthetic oligos were designed, each having a 30 base pair segment deleted, ablating any motifs within that particular segment. These ablations were designed to query the differential enhancer activities of 150bp oligos with and without the structured DNA sequence content of the 30 base pair subregion in question. We refer to the collection of wild-type, and non-wild-type oligos drawn from a single 150 base pair locus as tile sets and the individual 150 base pair oligos as “tiles”, or simply “oligos”. Tiles replacing a given 30 base pair “query sequence” with random DNA are referred to as “ablating” that section of DNA, and putative motifs in the query sequence are referred to as “ablated”.

We performed STARR-seq on the library of designed oligo nucleotides in U2OS cells, which express APOBEC3B at a high level. We also performed the same experiment in HCT116 cell lines--which do not express APOBEC3B--in order to identify regulatory elements associated with differential expression of APOBEC3B. Each DNA molecule in our library was barcoded with a random barcode to quantify DNA library diversity and adjust read counts to the underlying occurrence of DNA molecules. We observed approximately 41 DNA barcodes per oligo in U2OS and HCT116 cells and ~80 DNA barcodes per molecule in DLD1 cells. In our STARR-seq experiment, expressed transcripts were also tagged with random molecular UMIs identifying unique RNA molecules. The combination of RNA UMIs and DNA barcodes was designed to allow us to correct for sequence amplification bias, and DNA library skew in order to correct numerical artifacts to compute STARR-seq expression levels of a given synthetic DNA molecule.

### Oligo library construction and quality filtering

We searched for each barcode in the transcript sequencing data and quantified the number of unique transcript UMIs associated with each barcode to assess transcription rate. Transcription rate was quantified in two replicates each of three cell types, U2OS, DLD1 and HCT116. Using the number of transcript UMIs, normalized by the number of DNA barcodes observed, we computed a normalized transcription rate, “μ”, for each wild-type and each mutated oligo. In our aggregate data pooled over all cell types, 5041642 barcodes and 11834557 total UMIs are observed, yielding 2.3 unique transcripts per barcode and on average 76 transcript UMIs per olgo unique DNA sequence. The number of transcripts was counted per DNA barcode, added together for all barcodes associated with a given DNA sequence tile, and divided by the number of barcodes associated with that DNA tile. This number gives a normalized transcription rate for each DNA sequence tile, which we call mu, (“μ”) the normalized STARR-seq expression rate of a given oligo. This normalized expression rate was highly replicable between experiments, with a Pearson R of 0.93 (p<.1e-10).

## RESULTS

We demonstrated first that our technique would confirm the biological activity in synthetic constructs of known motifs corresponding to a transcription factor, NFKB1 previously suspected to activate the APOBEC-3B gene. Secondly, we sought to demonstrate that our expression-based perturbative analysis could resolve motif-containing ablations of highly expressed oligos which were enriched for motif instances of an activator an activator AP1 and a repressor PLAG1 with previously un factors may have biological significance in our understanding of cancer biology. Finally, we performed a broad regulatome-wide analysis of the activity levels of 579 motifs in the JASPAR motif database.

### Transcription activation by a known motif for the NFKB-1 transcription factor

Using 354 motif-containing instances and 122 motif-ablating instances in two replicates of U2OS cells, we observed that the NFKB motif positively enhances transcription. To assess the transcription activation potential of DNA sequence motifs thought to bind the NFKB-1 transcription factor, we computed normalized expression levels for all library oligos in our study. Expression levels were characterized and oligos were grouped by expression level, with the 95th-percentile expression level and 99.5th expression levels computed. (Fig 1a). It was observed that the top 99.5% of oligos by expression frequently contained a DNA binding motif for the NFKB-1 transcription factor, with the vast majority of high expression oligos across two replicates in U2OS cells, containing the NFKB1 sequence logo.

**Figure 1.**
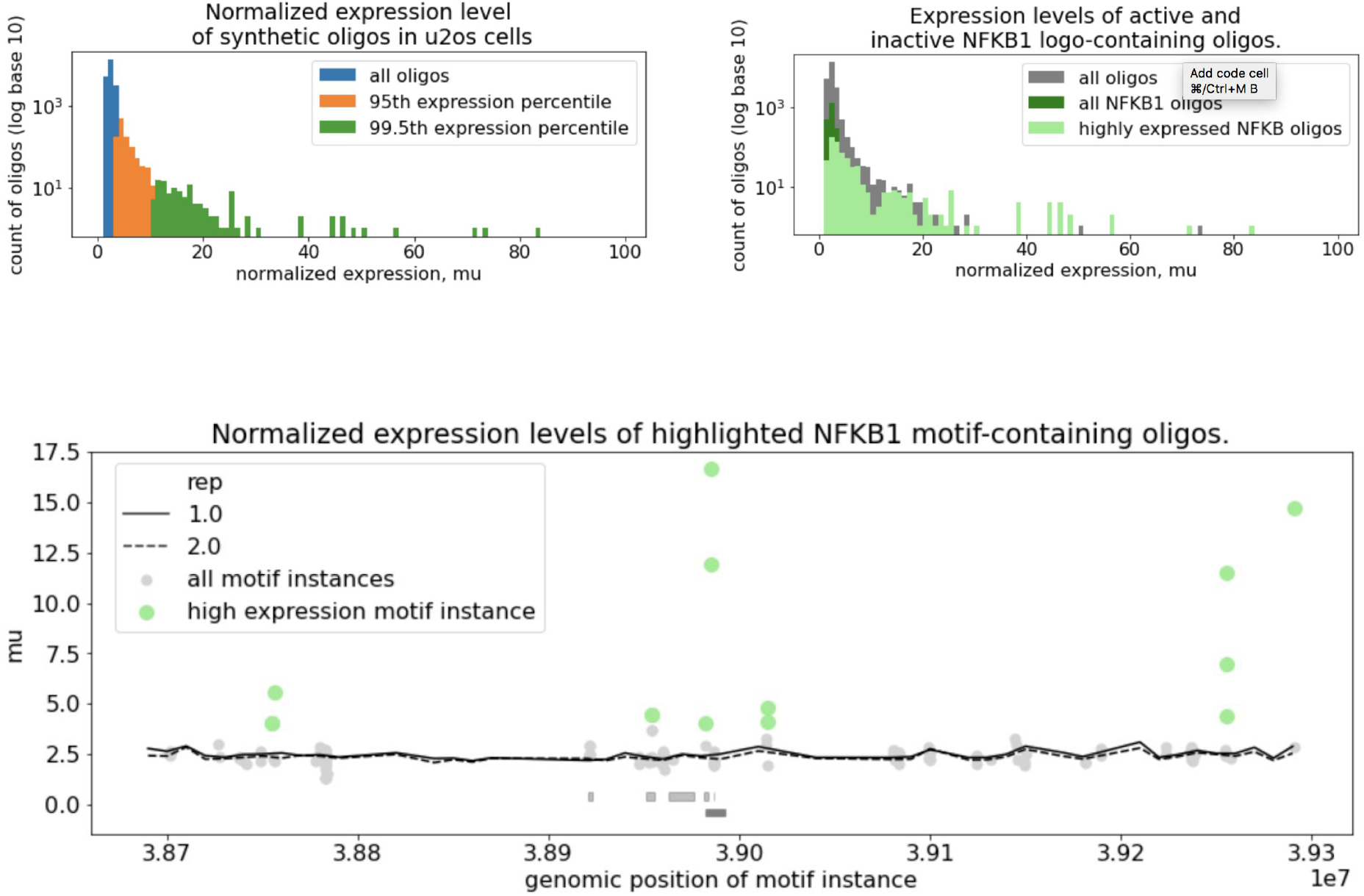
Expression levels of highly expressed synthetic oligos and occurrence of NFKB1 Motifs. Fig1(a), a histogram was computed of normalized expression level in order to identify highly enriched oligos. We computed 95th and 99.5th percentiles of synthetic oligos, looking at expression levels of STARR-seq synthetic DNA sequences in U2OS cells. Fig1(b), highly expressed synthetic oligos in U2OS cells were found to be enriched for NFKB1 motif instances. By using a fuzzy string matching approach with the NFKB1 sequence motif logo, we identified putative NFKB1 motif instances in the majority of high- and very high-expression synthetic oligos. Fig1(c), NFKB1 motif-containing oligos were mapped in the genomic vicinity of the APOBEC3B region. Points indicated in green represent expression levels of oligos containing the NFKB1 motif.

Expression levels of NFKB-containing oligos were averaged over two replicates in U2OS cells and plotted in Fig1c as a scatter plot over the APOBEC3B gene region, and apparently active enhancers were found in vicinity of the gene and a suspected cis-activating enhancer in addition to distal DNA sequence regions up to 300kb away from the APOBEC-3B gene.

14 Instances of the NFKB1 motif were studied in the 600kb genomic region surrounding the APOBEC3B gene. In (Fig2a), all tile sets overlapping each instance, tiles were split into motif-ablating and motif-intact and expression levels were compared with a violin plot. Shown in orange and blue is a histogram having raw counts of all tile instances generated from wild-type tile sets originally containing the NFKB1 binding motif. Raw counts in blue are expression levels of wild-type oligos, and non-wild-type oligos having benign 30bp mutations not affecting the NFKB1 binding motif. Raw counts in orange are expression levels of 30bp mutants generated from wild-type oligos containing the NFKB motif, but with synthetic ablations disrupting the motif. In (Fig2b) expression levels are shown for all instances and the Kolgomorov Smirnov test of the independent means is computed yielding a significant p-value=1.48e-7.

**Figure 2.**
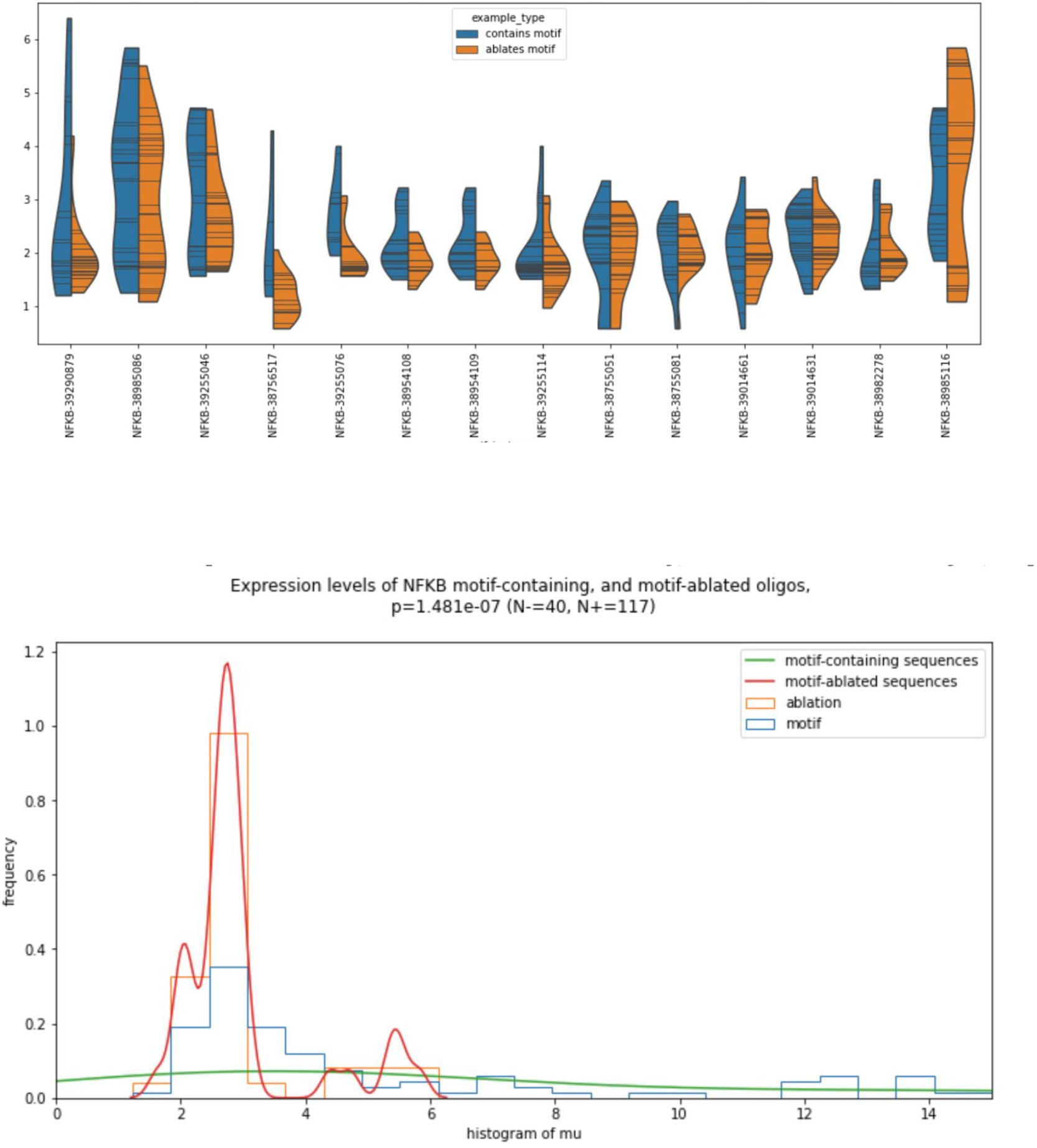
Expression Statistics of NFKB1 Motif Instances. Fig 2a. Log 2-fold differential expression levels of 14 NFKB1 motif instances in the search region Fig 2b. Expression levels of all NFKB1 motif-containing and ablated oligos across all motif instances.

For all library elements in each APOBEC3B tile set across two replicates in U2OS cells, each replicate of each tile was labeled accordion to whether the synthetic oligo contained the NFKB1 motif sequence or whether the APOBEC3B sequence was lost due to a 30bp synthetic ablation in that tile. Expression levels of all tiles in both replicates for U2OS cells are shown in Fig 1d, using the log-2 expression levels of the motif-containing and motif-ablated oligos shown as two separate populations.

To determine the likelihood that the NFKB1 motif was a factor in determining the normalized expression levels of STARR-seq oligos, we pooled the data across both replicates in our study of our library and compared the population of oligos having intact NFKB1 motifs with all oligos in the NFKB1-containing tile sets, but with NFKB1-containing query sequences ablated (Fig2b). Performing a Kolmogorov–Smirnov test to determine whether the 50 ablated examples and 117 intact examples were drawn from two different distributions, we found with a highly significant p-value of approximately 1.5*10^-7 indicating that the distributions of expression levels observed for ablated vs intact oligos were very likely different.

The NFKB1 motif can be confidently asserted to have an effect on the expression levels of synthetic oligos in U2OS cells.

### The AP-1 DNA sequence motif is correlated with enhancer mediated mRNA expression

We sought to extend the results of our investigation from the motifs of NFKB1 to transcription factor motifs to explore the possibility of their regulatory activity in U2OS cells. We examined four transcription factor motifs found in highly-expressed oligos, NR1, ESR2, and AP-1, in addition to NFKB1. For each of these transcription factor motifs, we hand-curated an ambiguous sequence logo using the highly expressed sequences and logo information in the JASPAR database (Figure 3a). We then searched for these sequence logos using an Acora-based4 string search in with a fuzzy string matching database allowing us to probe sequences within a hamming radius of our curated motif. Motif locations within a hamming radius of one are shown in Fig 3b.

**Figure 3.**
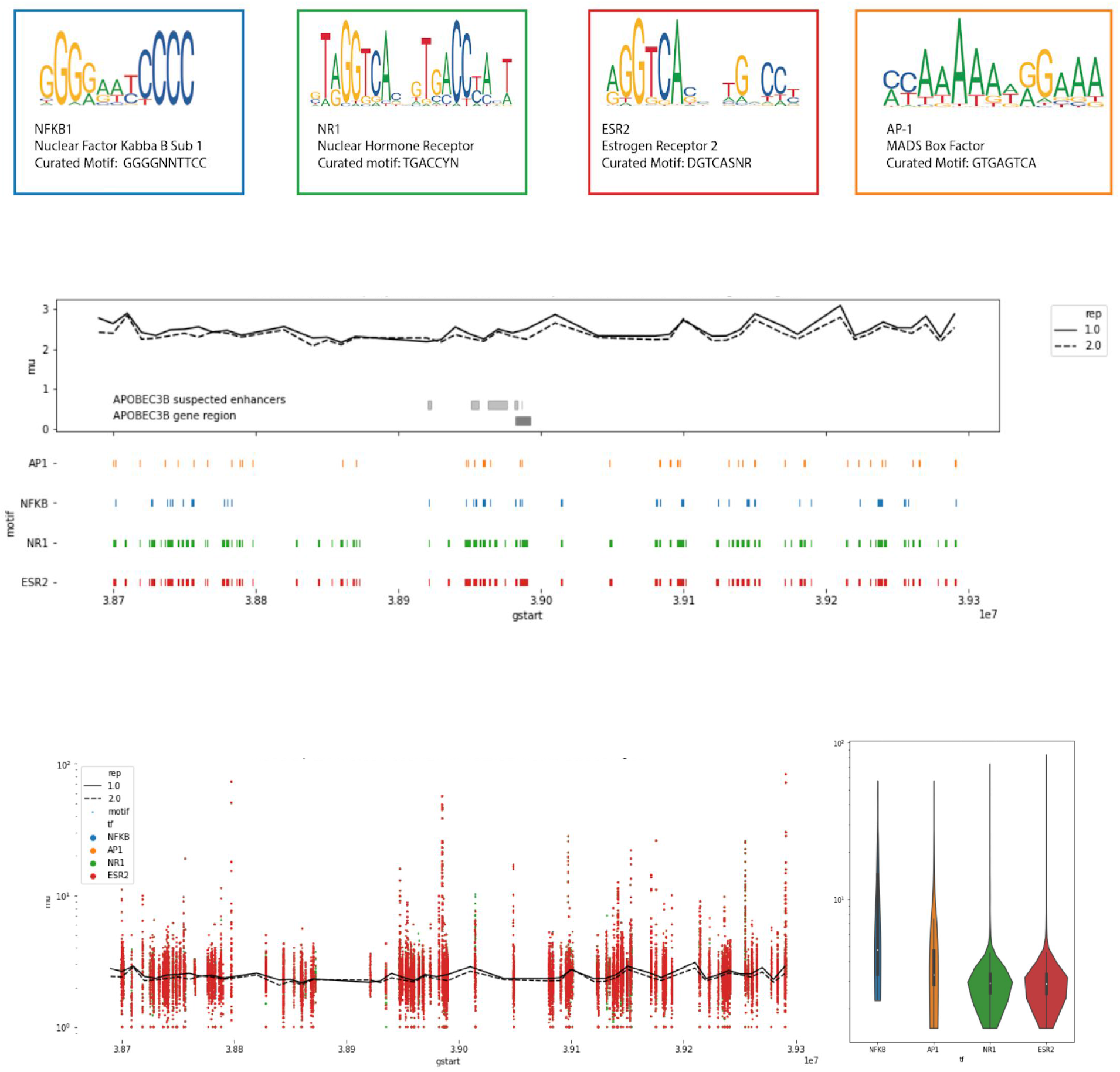
Investigation of hand-curated high-expression motifs. Fig 3a. Curated Motif Set sequence logos. Fig 3b. Curated Motif Set instance locations in APOBEC3B region Fig 3c. Expression levels of Curated Motif Set in APOBEC3B region Analysis of motif instances of hand-curated high-expression motif set. Hand-curated sequence logos for NR1, ESR2, AP1 and NFKB1 were defined using the JASPAR databases and motif observations in the high-expression synthetic oligos (Fig3a). The synthetic oligo library was scanned for all instances of the hand-curated motif set, using an added hamming search radius of +1 in the APOBEC3B region to allow fuzzy matching (Fig3b). Expression levels for each motif were plotted on a spatial axis, and combined into a violin plot showing the aggregate expression levels of instances of each motif. (Fig3c)

For each of these four putative transcription factor motifs, we studied the distribution of expression levels of all oligos in our database to understand the differential expression of oligos containing the putative motif versus oligos lacking the motif due to a 30 base pair synthetic ablation but otherwise sharing DNA context.

For NFKB1 and AP-1 motifs, significant changes in expression (p-values of 1.5×10^-7 and 5*10^-4, respectively) were found by comparing expression levels of oligos containing motif sequence logos with otherwise similar oligos having 30 base pair ablations disrupting those oligos, (Fig4a) For exact logo matches and for fuzzy logo matches within a hamming radius of 1 (Fig4b), both the NFKB and AP1 motifs appeared to impart a significant change in expression levels, suggesting that instances of these sequence logos imparted a significant change in expression levels of exogenous DNA in the STARR-seq assay.

**Figure 4.**
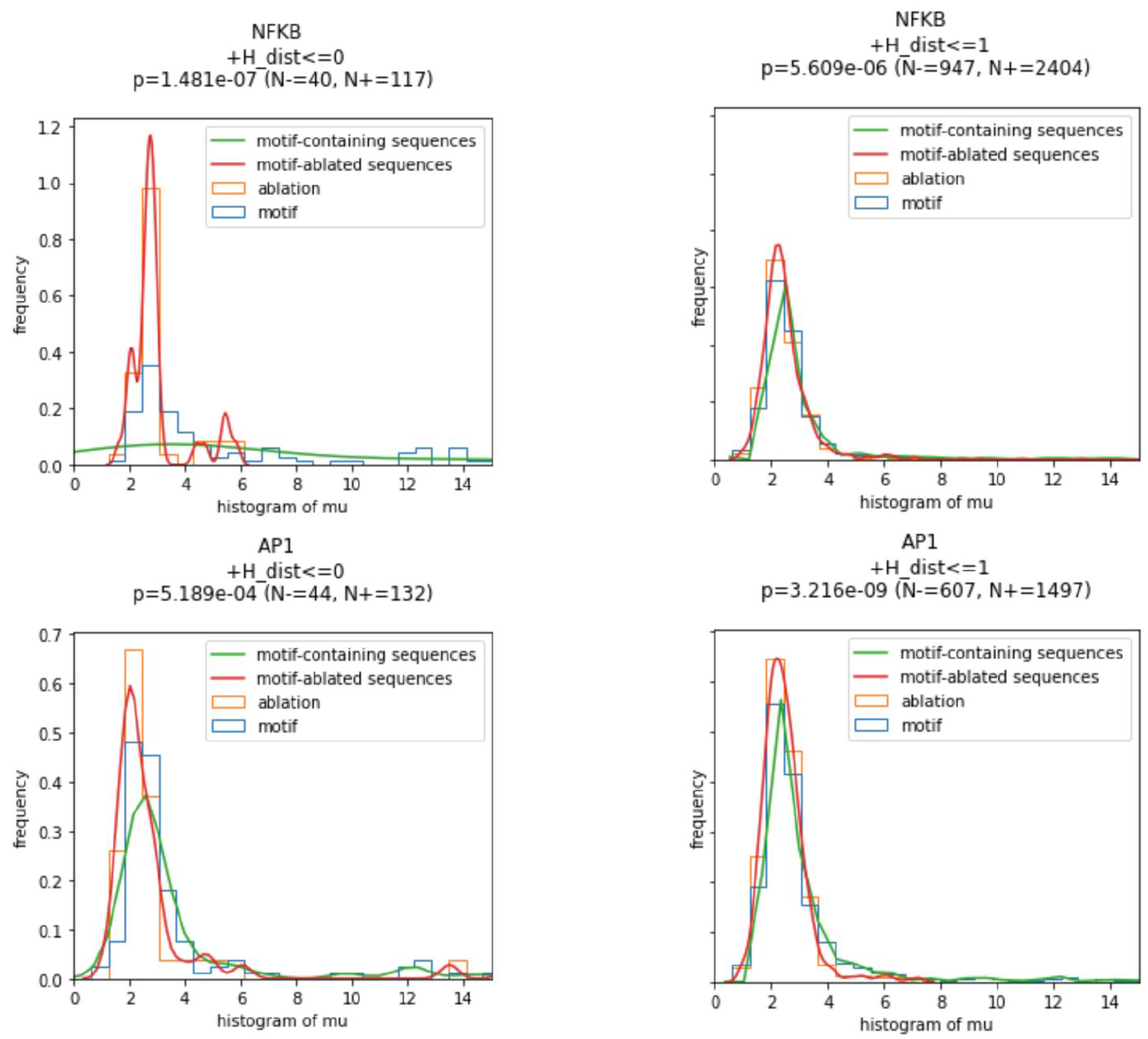
Differential distributions of expression levels for motif-containing, and motif-ablated synthetic oligos in U2OS cells for NFKB1 and AP-1 motifs. Fig 4a: Significant changes in expression level of NFKB & AP1 sequence-containing and motif sequence-ablating oligos. Exact logo matches. Fig 4b: Significant change in expression level of NFKB & AP1 sequence-containing and motif sequence-ablating oligos. Max hamming distance 1 logo matches.

### Potent Repressor And Activator Motifs are identified from 579 candidates

We applied STARR-scan to a curated database of 579 motifs in the JASPAR database to form a more complete picture of the regulatory potential of DNA sequences in the APOBEC-3B region in U2OS cells. For each motif, we scanned its canonical motif position weight matrix provided in the JASPAR database over all oligo DNA sequencesin our study and scored each oligo for motif presence or absence. For each transcription factor motif, we computed the mean expression level of oligos containing that motif and oligos with matching context but the motif ablated.

We observed that the ablation of some transcription factor motifs appeared to substantially shift oligo related mRNA expression levels, and thus we sought to identify the top repressive and activating motifs from the JASPAR database. We chose three motifs that caused the greatest increase and decrease in mRNA expression as candidate activator and repressor motifs respectively.

Amongst the potential repressor motifs, TCF7L1 was the most prevalent. TCF7L1 motifs occurred at five distinct positions in our dataset. Putative repressor element SP4 occurred in possible cis regulation sequence near the APOBEC-3B promoter and at a second location in the putative distal enhancer region. Putative repressor MEF2D motif occurred at only one position, and putative repressor TCF7L1 motif appeared in five loci throughout the region under study. FOS / JUN and JDP2 motifs appeared in cis-regulating enhancer and FSL2:JUNB occurred solely in distal regions remote from the APOBEC3B gene.

A search was performed for all instances in our dataset of 579 motifs in the JASPAR database. 503 motifs were found in at least one oligo in our library. For each motif, a mean expression level was computed for all wild-type oligos containing that motif and a second mean expression was computed for those oligos having a 30-bp ablation which eliminate that motif instance but kept intact the surrounding context. Expression levels of wild-type and ablation-containing oligos are shown in a histogram in (Fig5a). For each of the 503 motifs with at least one occurrence in our wild-type library, the difference in mean expression levels between wild-type and motif-ablating oligos was computed and shown in a histogram, (Fig5b). The .5% of motifs whose elimination caused the greatest negative change in expression level, FOSL2, JDP2, and FOSB are possible activator motifs, and the .5% of motifs whose elimination caused the greatest increase in expression level, SP4, TCF7L1, and MEF2D are putative repressor motifs.

**Figure 5.**
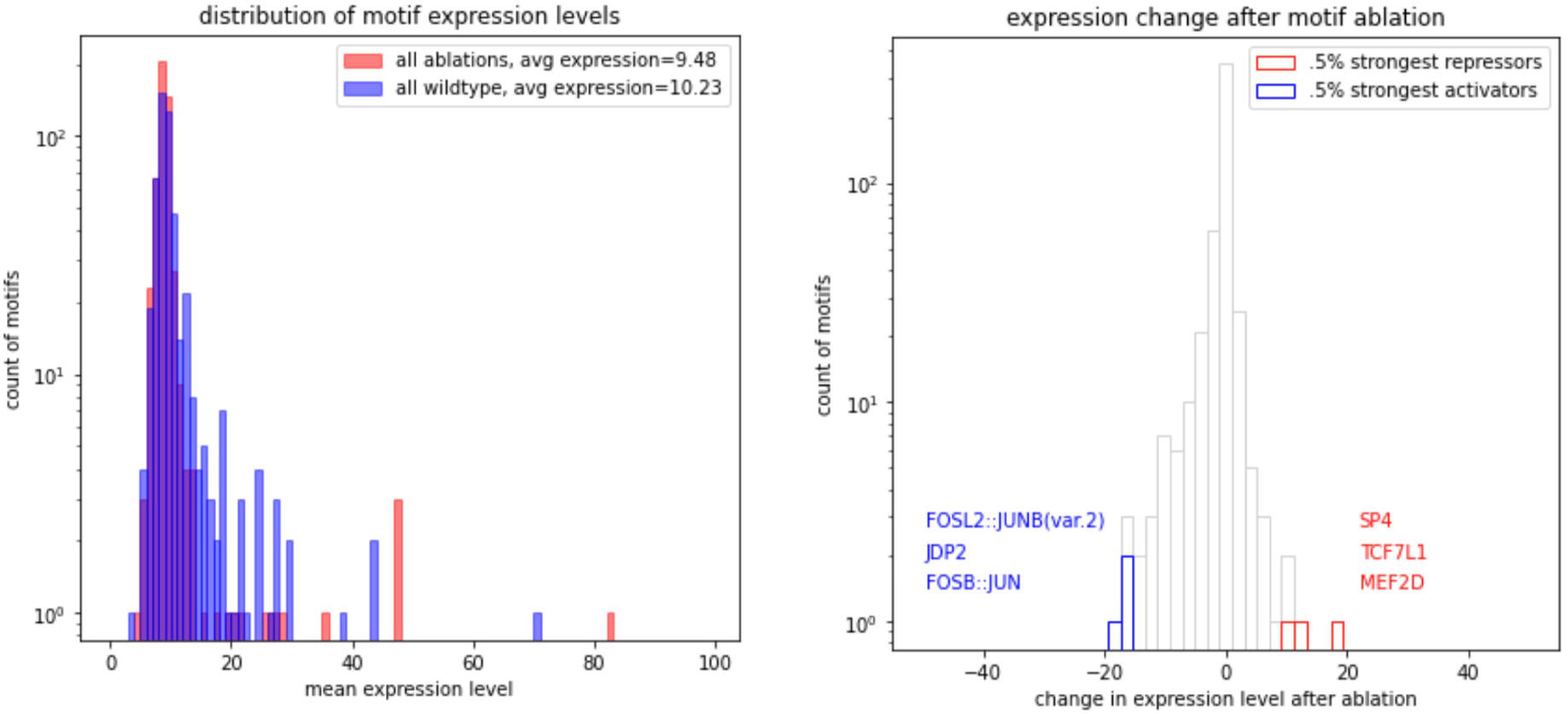
Expression levels of 579 JASPAR TF motifs and context-preserving synthetic ablations. Fig 5(a) Expression level of 579 intact JASPAR TFs and their ablated variants. Fig 5(b) Expression level change following ablation for 579 JASPAR oligos.

**Figure 6.**
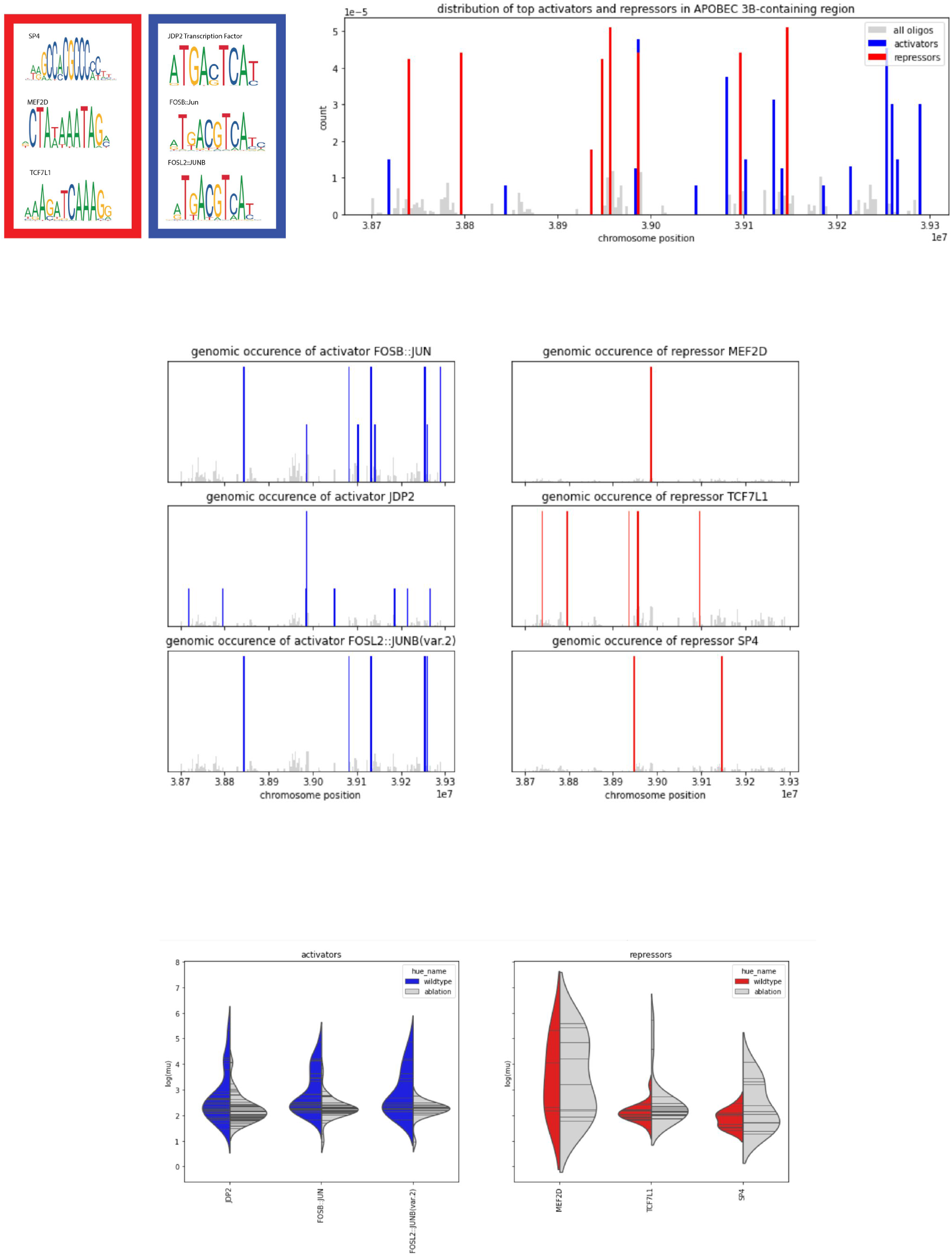
Spatial distribution and expression levels of activator and repressor motifs. Fig 6a: PWMs and overall spatial distribution of all activators and repressors in the region under study. Fig 6b: Individual instances of each repressor and activator DNA motif were identified in the genomic surrounding APOBEC3B with frequencies plotted in a histogram. Fig 6c: Distribution of expression levels for motif-containing vs motif-ablated oligos for top activators and repressors were computed, and the distributions show for each putative activator and enhancer.

Encouraged by these results, we sought to perform a statistical analysis for all transcription factor motifs imparting statistically confident changes in expression levels in U2OS cells. To perform this analysis, we performed the analysis of Fig4c for 503 of 579 JASPAR motifs with at least one instance in our library, and on each motif performed a Kolgomorov Smirnov 2 sample test on all data points, keeping two replicates separate in U2OS cells. For each transcription factor, we tested the hypothesis that intact sequence motifs changed the STARR-seq normalized expression level mu in each replicate of our experiment in U2OS cells. (Fig7) Statistical tests were carried out to identify transcription factor motifs causing a highly significant change in expression (Fig7). 86 transcription factors from the JASPAR database had a significant p-value <.05 after combination of replicates with Fisher’s method and multiple hypothesis correction using the bonferroni method with n=579. Using Fisher’s method to combine two replicates of our experiment in U2OS cells, our identified 25 motif candidates which appear to effect expression levels with very high confidence, p<5e-7 (Fig8a), and found a significant correlation (R=.64) between motif significance measurements in U2OS and HCT116 cells (Fig8b).

**Figure 7.**
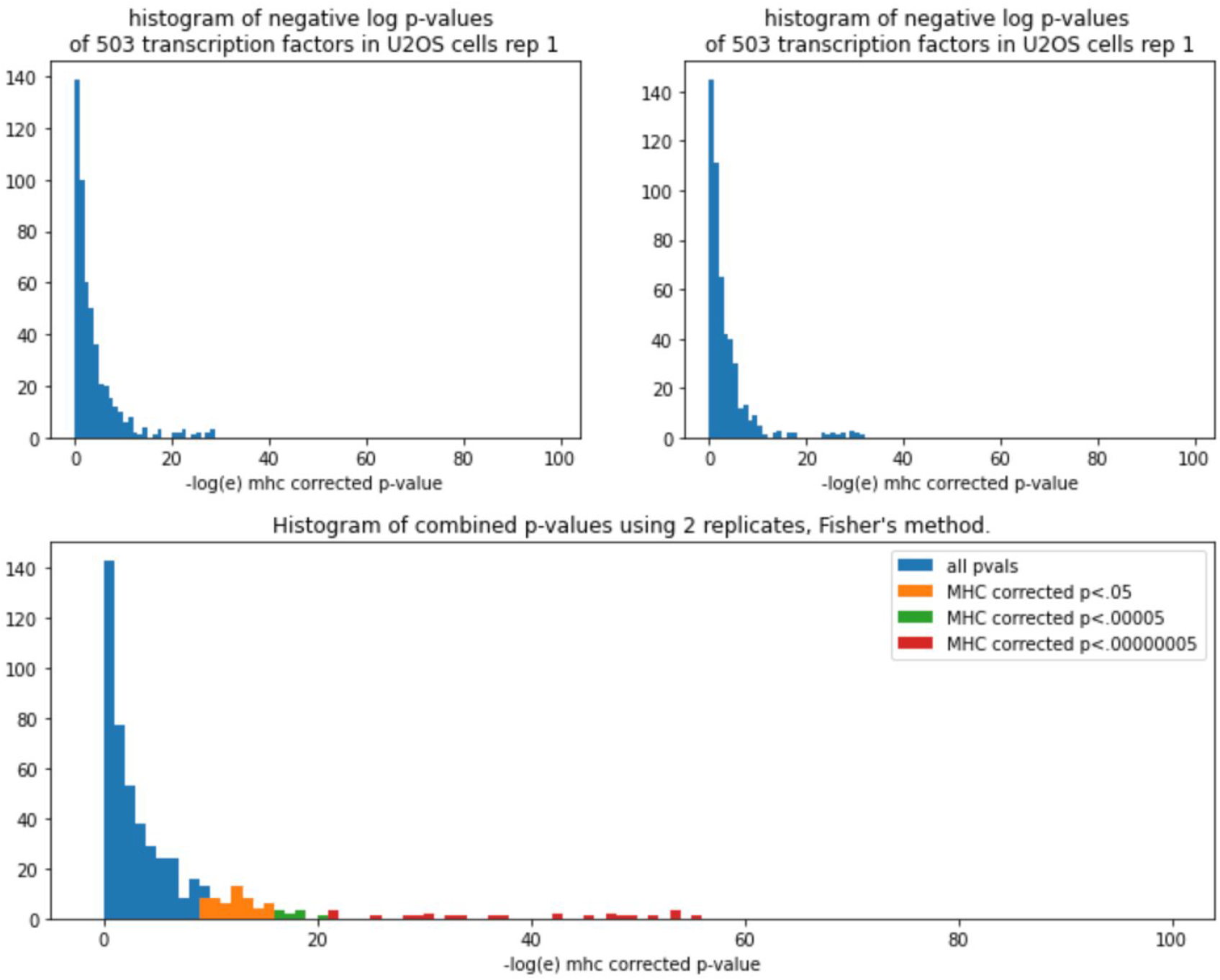
Statistical tests of the likelihood of regulation. Fig7. A comprehensive screen was performed by identifying motif instances for 579 annotated transcription factor motifs in the JASPAR database. Scanning for motif containing tile sets for each motif in the database, we split the synthetic oligos into populations of tiles having intact motif sequences and populations having those motifs ablated. Performing a non-parametric Kolgomorov Smirnov test on all instances of each motif in two replicates, we were able to assess the p-value that any transcription factor had regulatory activity in each replicate. Combining P-values with Fisher’s method we were able to compute a p-value using the replicate data. 86 transcription factors appeared to have a significant effect, (p<.05,bonferroni multiple hypothesis correction, n=579), and even taking a very stringent p-value of 5*10^-7, 24 transcription factors passed our test including a number of factors suspected to activate APOBEC3B. Highly significant transcription factors tended to have high values of average normalized expression, mu, which was computed in HCT116 and U2OS cell lines.

**Figure 8.**
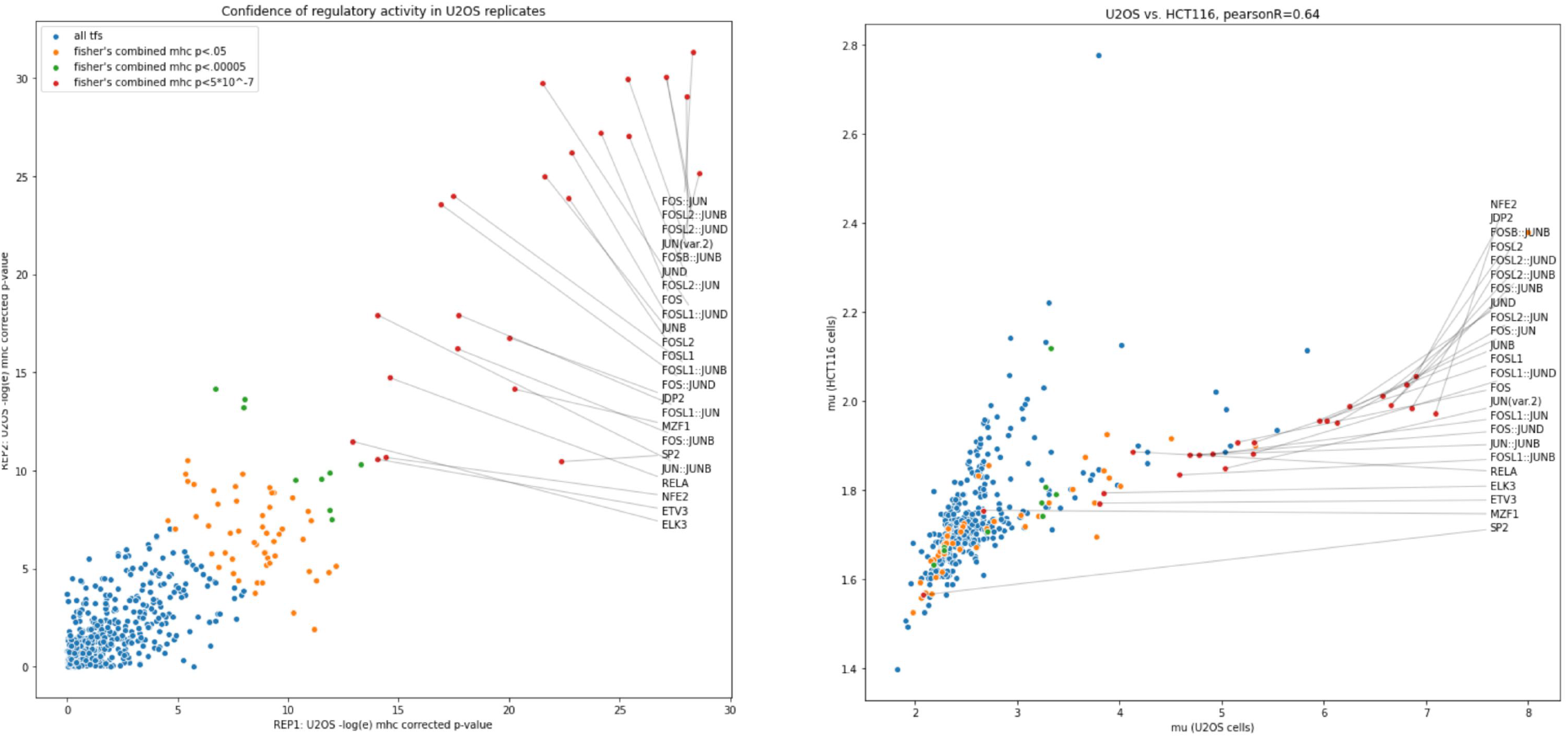
Replicability of significant regulating motifs. Fig8a: Combined P-values from two replicates. Indicated in red are highly significant transcription factors p<5*10^-7, computed by Fisher’s method, combining replicates of two STARR-seq experiments in U2OS used in this study. Fig 8b: Expression levels of significant transcription factors in U2OS, HCT116 cell lines. A strong correlation (R=.64) is found between HCT116 and U2OS cell lines.

## DISCUSSION

We demonstrated up-regulated reporter expression caused by 150 base pair synthetic enhancers from genomic regions of interest. We found that within each of these regions it was possible to identify smaller regulatory subregions with a tiled ablation strategy, with enhancer sequence ablations causing replicable and cell type-specific expression level changes. Using a database of known position weight matrices for characterized transcription factors, including factors known to regulate APOBEC3B, it was possible to identify which motifs were regulatory in cellular conditions in each sample.

We presented a general strategy for discovering the regulatory sequences in tiled STARR expression oligos. This strategy can be applied to putative enhancers in other regions of interest to effectively characterize sequences important for gene regulation. We quantified the regulatory effects of individual motifs by characterizing the significance of observed expression changes between wildtype and ablated enhancer sequences.

In addition to presenting a general strategy to scan genomic regions for regulatory DNA at high resolution, our work also provided regulatory results specific fo APOBEC3B. We confirmed the activity of NFKB / RELA in U2OS cell lines which express APOBEC3B, this work confirms past work indicating that NFKB1 may have a role in regulating the expression of that gene.

Furthermore our analysis revealed at least one apparent repressor motif, SP2 from our unbiased screen using the JASPAR database. SP2 has previously been suggested to have a role in cancers, with SP2 having been possibly indicated as a transcriptional repressor of antigen-related cell adhesion molecules ^5^. Furthermore, we identified the motifs of activators which could be related to the regulation of APOBEC3B, including MZF1 which is suspected to have a role in cancer metastasis ^6, 7^, NFE2 ^8^, and JDP2^9^, in addition to a number of FOS / JUN family transcription factors. This study strongly suggests that DNA motifs previously linked with these transcription factors may have regulatory significance in the APOBEC3B gene region and in the case of some, the activity may have cell-type specificity with differential levels of activity between HCT116 cells and U2OS cells.

## FUTURE WORK

Recent work has explored the regulatory potential of single transcription factors and the combinatorial logic of multiple factors in specific cell types^3^. Future work could build upon our methods to explain the combinatorial regulation of multiple motifs using a non-binary scoring process. For example, logistic regression or a neural network could be used to predict expression levels from sequence features of oligos. The interpretation of these models could be used to assess the individual and combinatorial contributions of motifs in addition to sufficiency of single or combined motif instances suggesting factor interactions such as that suggested between REL and NFKB1 in this studied in a systematic fashion.

The STARR-scan methodology can precisely identify regulatory motifs, which we expect will permit it to characterize the effect of single nucleotide variants on gene expression ^10^ The improved analysis of genetic variation will permit us to understand the genetic underpinnings of disease, and propose new therapeutic interventions.

## REFERENCES

1. Mathur, D. et al. Analysis of the mouse embryonic stem cell regulatory networks obtained by ChIP-chip and ChIP-PET. Genome Biol. 9, R126 (2008).

2. Rajagopal, N. et al. High-throughput mapping of regulatory DNA. Nat. Biotechnol. 34, 167–174 (2016).

3. Guo, Y., Mahony, S. & Gifford, D. K. High resolution genome wide binding event finding and motif discovery reveals transcription factor spatial binding constraints. PLoS Comput. Biol. 8, e1002638 (2012).

4. Aho, A. V. & Corasick, M. J. Efficient string matching: an aid to bibliographic search. Commun. ACM 18, 333–340 (1975).

5. Phan, D. et al. Identification of Sp2 as a transcriptional repressor of carcinoembryonic antigen-related cell adhesion molecule 1 in tumorigenesis. Cancer Res. 64, 3072–3078 (2004).

6. Brix, D. M., Bundgaard Clemmensen, K. K. & Kallunki, T. Zinc Finger Transcription Factor MZF1-A Specific Regulator of Cancer Invasion. Cells 9, (2020).

7. Mudduluru, G., Vajkoczy, P. & Allgayer, H. Myeloid zinc finger 1 induces migration, invasion, and in vivo metastasis through Axl gene expression in solid cancer. Mol. Cancer Res. 8, 159–169 (2010).

8. Qian, Z. et al. Nuclear factor, erythroid 2-like 2-associated molecular signature predicts lung cancer survival. Sci. Rep. 5, 16889 (2015).

9. Bitton-Worms, K., Pikarsky, E. & Aronheim, A. The AP-1 repressor protein, JDP2, potentiates hepatocellular carcinoma in mice. Mol. Cancer 9, 54 (2010).

10. Zeng, H., Hashimoto, T., Kang, D. D. & Gifford, D. K. GERV: a statistical method for generative evaluation of regulatory variants for transcription factor binding. Bioinformatics 32, 490–496 (2016).

